# Characterization of spatial homogeneous regions in tissues with concordex

**DOI:** 10.1101/2023.06.28.546949

**Authors:** Kayla C. Jackson, A. Sina Booeshaghi, Ángel Gálvez-Merchán, Lambda Moses, Tara Chari, Alexandra Kim, Orhan Hosten-Mittermaier, Lior Pachter

## Abstract

The rapid advancement of spatially resolved transcriptomics (SRT) technologies has facilitated exploration of how gene expression varies across tissues. However, identifying spatially variable genes remains challenging due to confounding variation introduced by the spatial distribution of cell types. We introduce a new approach to identifying spatial domains that are homogeneous with respect to cell-type composition that facilitates the decomposition of gene expression patterns by cell-type and spatial variation. Our method, called concordex, is efficient and effective across technological platforms and tissue types, and using several biological datasets we show that it can be used to identify genes with subtle variation patterns that are missed when considering only cell-type variation, or spatial variation, alone. The concordex tool is freely available at https://github.com/pachterlab/concordexR.

## Introduction

Spatially resolved transcriptomics (SRT) have enabled highly multiplexed molecular profiling of cells within a tissue, with current technologies presenting a range of tradeoffs in approach and resolution (1). Broadly, in-situ hybridization based methods, such as seqFISH (2, 3), seqFISH+ (4), and MERFISH (5), offer cellular or sub-cellular resolution for capture of hundreds to thousands of genes, while methods that rely on spatial barcoding and sequencing (e.g. Visium, Slide-Seq (6), Slide-SeqV2 (7) offer near-cellular resolution and measure the expression of genes across the entire transcriptome.

A major goal of spatial transcriptomics data analysis is the determination of spatially variable genes. Ideally, methods should be able to distinguish whether variability is driven by the spatial distribution of cell types or by spatial variation that is independent of cell-type. One approach to untangling these two covariates is to partition assayed tissues into regions that constitute domains of functional or compositional homogeneity. This task first relies on abstracting transcriptomic expression into notions of cell type, whereby cells of the same type have similar transcriptomic profiles, but can be morphologically or functionally distinct. The concept of a spatial region introduces another layer of abstraction and requires aggregation of cell types into domains with distinct cell-type composition. The cells in these regions are characterized by their local cellular environments, and have neighborhoods with similar proportions of cell types, which can be a mixture of cell types or a single type. We therefore refer to regions with this property as spatial homogeneous regions (SHRs).

Several algorithms have been proposed for identifying spatial or tissue domains defined by coherent gene expression, yet, in principle, these domains can consist of various cell types with different expression profiles (8–13). The result is that these methods implicitly identify SHRs. Broadly, these approaches rely on neural networks, hidden Markov random fields (HMRFs), or spatial smoothing to encode spatial dependence. For example, Giotto (11) and BayesSpace (13) infer domain assignment using an HMRF and relate the gene expression of a cell or spot and its neighbors. BANKSY (8) uses spatial kernels to encode spatial dependence in the local and extended environment around a tissue. GASTON (12) relies on a neural network to represent gene expression and spatial information as a one-dimensional gradient. SpaGCN (10) and STAGATE (9) use graph convolutional neural networks to integrate gene expression with spatial and/or histology information.

The regions identified by these methods have been used in downstream analysis pipelines to detect spatial differentially expressed genes (sp-DEGs), but it is unclear how this analysis partitions variability into cell type and spatial effects. In an extreme case, a SHR composed entirely of a unique cell type—absent from other SHRs—may yield differentially expressed genes that simply reflect cell-type differences rather than true spatial variation. On the other hand, testing for domain differences without accounting for the effect of cell type may obscure distinct spatial effects and ignores cases where cell type and spatial effects overlap. Thus, the question of how to best identify SHRs and sp-DEGs in a coherent manner remains open.

We propose to solve these problems by first utilizing spatial k-nearest-neighbor (kNN) graph representation of transcriptomics data to answer questions about spatial homogeneity in SRT data. The key to our approach is a method we develop for assessing the neighborhood composition of nodes in a kNN graph built from spatial or non-spatial attributes, which we implement in a tool called concordex. We show that concordex can efficiently and effectively identify SHRs in spatial transcriptomics data, and also that it is a useful tool for assessing concordance between partitions of cells derived from clustering and kNN graphs in non-spatial transcriptomics data. We demonstrate the utility of concordex in many contexts with both simulated and publicly available biological datasets that encompass a range of technologies. Subsequently, we demonstrate the compatibility of concordex with a method for differential analysis based on a generalized linear model (GLM). We model SHRs and cell types directly, which allows for the separation of gene expression patterns driven by spatial context from those driven by cell type. This coupling of concordex to a GLM reveals genes with subtle, yet interesting, spatial variation patterns.

## Results

### Neighborhood consolidation with concordex

The con-cordex workflow can be used to interrogate the neighborhood composition of the nodes in a spatial kNN graph, *G* = (*V, E*), where *V* is a set of cells or spots and *E* is the set of edges in the graph. The edges of the graph are determined by some metric on *V*, usually by computation of transcriptomic or spatial distance, and the nodes are assigned predetermined discrete or continuous labels. When discrete labels are available, the concordex assessment proceeds first by calculating the neighborhood consolidation matrix *K*, with a row for each cell *i* and one column for each label *j* (Figure 1). The entries *K*_*ij*_ can be interpreted as the fraction of neighbors of cell *i* that are assigned label *j*. This representation has been used to identify ‘cellular neighborhoods’ in multiplexed imaging data (14–16), but has yet to be comprehensively applied to datasets on scale with modern SRT technologies. Additionally, since this approach relies on pre-annotated labels, which are often unavailable or imprecisely defined, concordex extends the labeling framework to include continuous representations of cells. Here, the columns of the neighborhood consolidation matrix correspond to each component of the continuous vector. Clustering the rows in the neighborhood consolidation matrix assembles cells into SHRs, where cells within an SHR can be thought of as having similar neighborhood composition. Though we focus on the spatial applications of concordex, the matrix *K* can be used in a non-spatial context to reveal cells with non-homogeneous neighborhoods and assess cluster boundaries via direct visualization of between-cluster relationships and within-cluster heterogeneity (Supplemental Note).

**Fig. 1.**
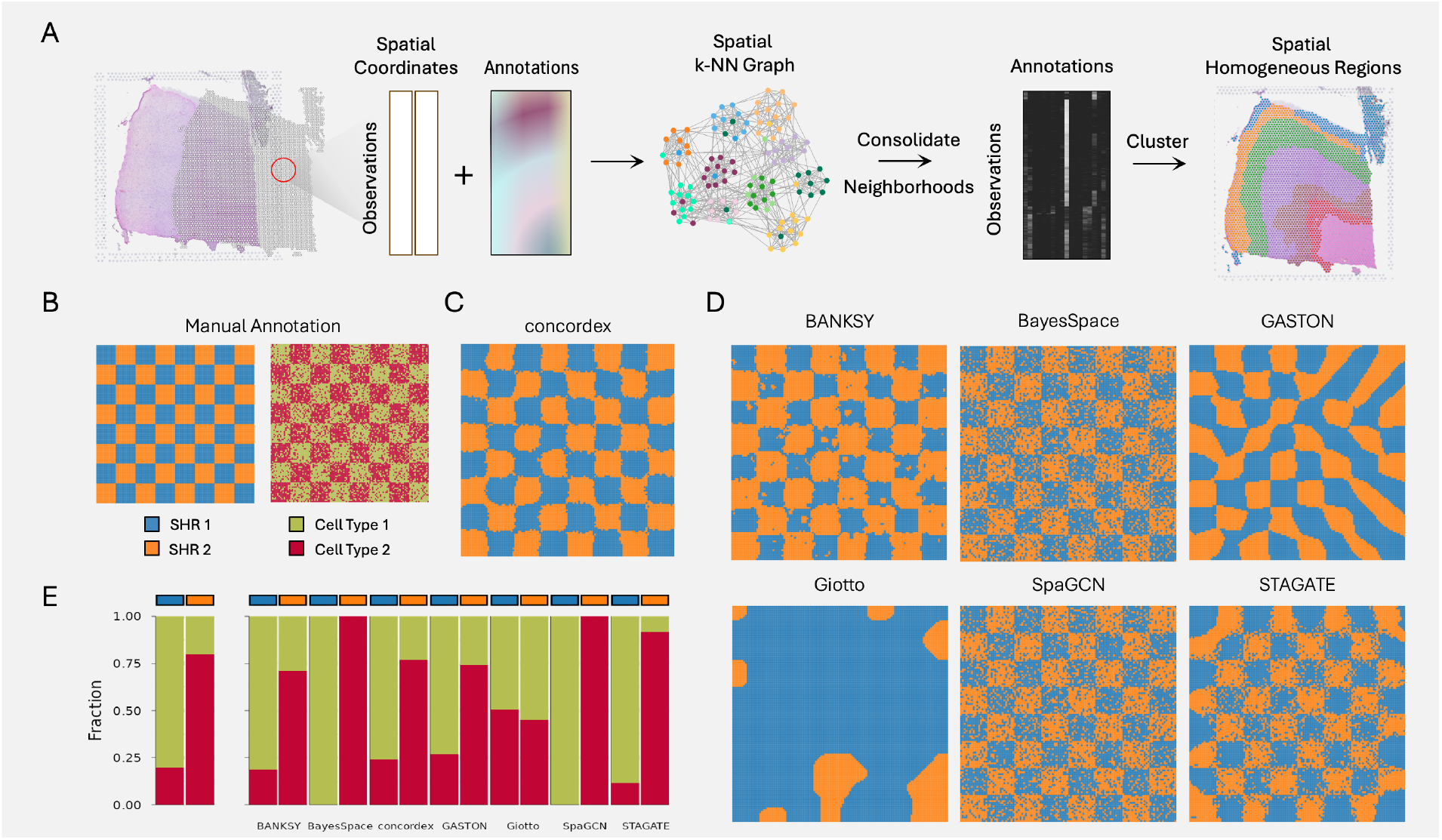
The concordex workflow: **A**. A spatial kNN graph is constructed from spatial coordinates and associated annotations such as cell types or features from a principal component analysis projection. For each cell or spot, the entries in the neighborhood consolidation matrix represent the fraction of neighbors that have a label indicated. SHRs are defined by clustering the neighborhood consolidation matrix. **B**. A control experiment in which a chessboard pattern consists of two regions, each comprising two cell types, one with 80% of one cell type, and 20% of another, and the other region with a 20% / 80% mix. **C**. concordex captured this pattern with high accuracy, while **D**. the performance of six other methods varied. **E**. The expected proportion of cell types in each SHR in the control (left) and the predicted proportion of cell types produced by each method (right)

We use concordex to identify SHRs in datasets encompassing several SRT technologies and spatial scales. We build on previous results that demonstrate the utility of this approach with discrete annotations and extend this result to show that continuous labels accurately partition cells into SHRs. In cases where anatomical information about the tissue is known, we use k-means clustering to identify SHRs in the spatial context, but otherwise, SHRs are obtained using graph-based clustering algorithms such as Leiden (17) or Louvain (18) to cluster the neighborhood consolidation matrix.

### Benchmarking concordex in control and Visium data

To better understand the utility of our approach, we simulated control spatial datasets in various patterns. First, we designed a synthetic dataset containing two cell types and distributed the cells on a chessboard in different proportions (Figure 1B, Methods). This scenario is useful because it allows analysis of whether a method can detect regions of varying cell type composition, even when a cell type is present throughout the entire spatial field of view. We assessed whether other methods described above could perform the same region segmentation task. Ideally, methods should detect the checkerboard as a macro-pattern rather than the exact positions of the individual cell types.

We find that concordex is able to effectively reconstruct the chessboard squares (Figure1C) and each detected region contains the expected proportion of the simulated cell types. The concordex predictions were most similar to STAGATE and BANKSY, with both methods producing recognizable chessboard and correctly assigning grid points with relatively high accuracy (Figure 1D). Conversely, other methods failed to perform this task in notable ways. Two methods, BayesSpace and SpaGCN, reproduced the cell type assignment rather than aggregating the points into regions even when using parameters that should prefer region identification over cell type identification (Figure 1D). Though the cell type locations resemble the chessboard grid, we argue that the misidentification of regions at this step precludes meaningful downstream gene analysis. On the other hand, Giotto and GASTON did not produce a recognizable chessboard (Figure 1D). When we arranged the simulated cell types in sequential layers and used concordex to predict the layers (Supplemental Figure S6), we found the results were consistent with the chessboard control (Supplemental Figure S6). Importantly, concordex captures the expected organization of the simulated tissue across an array of gene expression patterns and relies on compositional changes, not expression, to determine regional boundaries.

Next, we evaluated the ability of concordex to identify SHRs in 12 manually annotated sections of the human dorsolateral prefrontal cortex (DLPFC) (19). This dataset was acquired on the 10x Genomics Visium platform, and each section includes up to 7 annotated anatomical regions that can serve as ground truth. Since the Visium platform does not have cellular resolution and deconvoluted cell type predictions were not available for the spots, we used the first 50 principal components to compute the neighborhood consolidation matrix. Given the presence of unbalanced clusters, we used the adjusted mutual information (AMI) to compare the accuracy of SHRs identified by concordex to established and recently developed spatial clustering methods, including BANKSY, BayesSpace, GASTON, Giotto, SpaGCN, and STAGATE.

Across the 12 sections, concordex consistently identified the expected cortical layers across sections of the DLPFC dataset (Figure 2A)). Using sample 151675 as an example, we found that the SHRs identified by concordex generally agree with the ground truth annotations (Figure 2B) and represent a qualitative and quantitative improvement over several methods (Figure 2C-D), confirming that the neighborhood consolidation matrix is a useful representation for defining SHRs in practical applications. The other methods produced results that vary in their agreement with the annotations (Figure 2D). Despite having comparable AMI (Figure 2A) in some sections, we find that the predictions from other methods produce layers that are qualitatively less distinct than concordex, and in general, require more computational time to compute (Supplmental Figure S4).

**Fig. 2.**
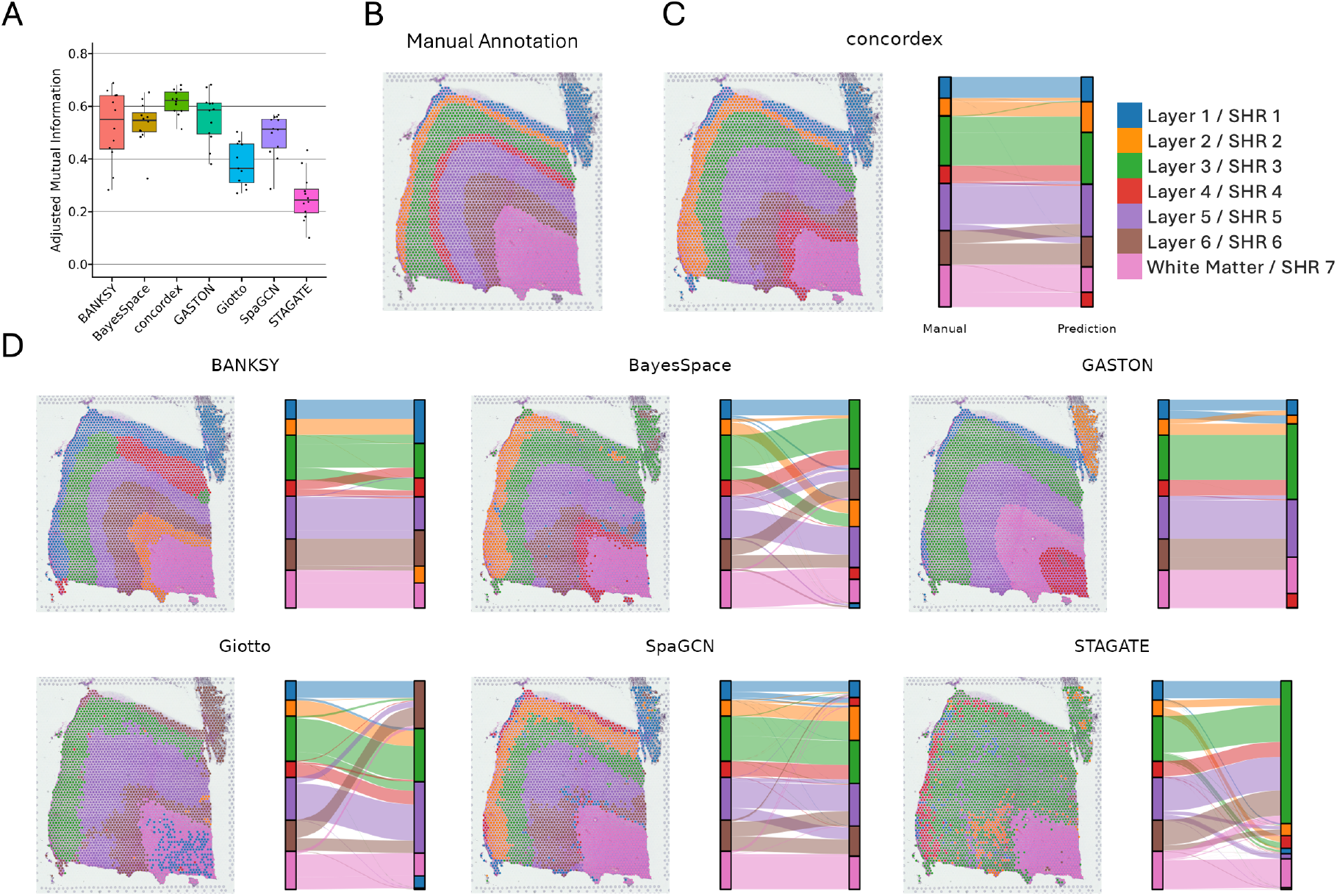
Evaluation of in Visium DLPFC dataset. **A**. Comparison of adjusted mutual information scores across 12 sections of the DLPFC dataset **B**. The manual annotations for section 151675. **C**. Evaluation of concordex and **D**. other methods in section 151675 from the human DLPFC dataset. Each method is shown next to the corresponding alluvial diagram mapping the manual annotations to the predicted annotations.

Interestingly, Giotto, SpaGCN, and STAGATE produced results that are substantially worse than those of the other methods and contrast with the results published in the respective manuscripts for these methods. This is in part due to the fact that we used analytic Pearson residuals (20) instead of sequencing depth to normalize the data before dimension reduction with PCA. Although the question of how to normalize spatial transcriptomics data remains open, we found that the PCs computed from analytic Pearson residuals better retained spatial tissue structure compared to explicitly depth normalized data.

### Improved identification of layers in the mouse neocortex using concordex

To assess whether concordex could be used to predict SHRs in a dataset with cellular resolution, we applied our method to an adult mouse primary visual neocortex profiled with STARmap technology (21). STARmap relies on padlock probes that hybridize to intracellular mRNA followed by in situ amplification and imaging to read out gene-specific sequences. The authors provided data for 1020 genes detected in 1207 cells along with molecularly defined cell types and annotated anatomical regions as shown in Figure 3A. To further demonstrate the utility of using continuous representation of cells, we again used the first 50 PCs as cell labels for input to concordex.

**Fig. 3.**
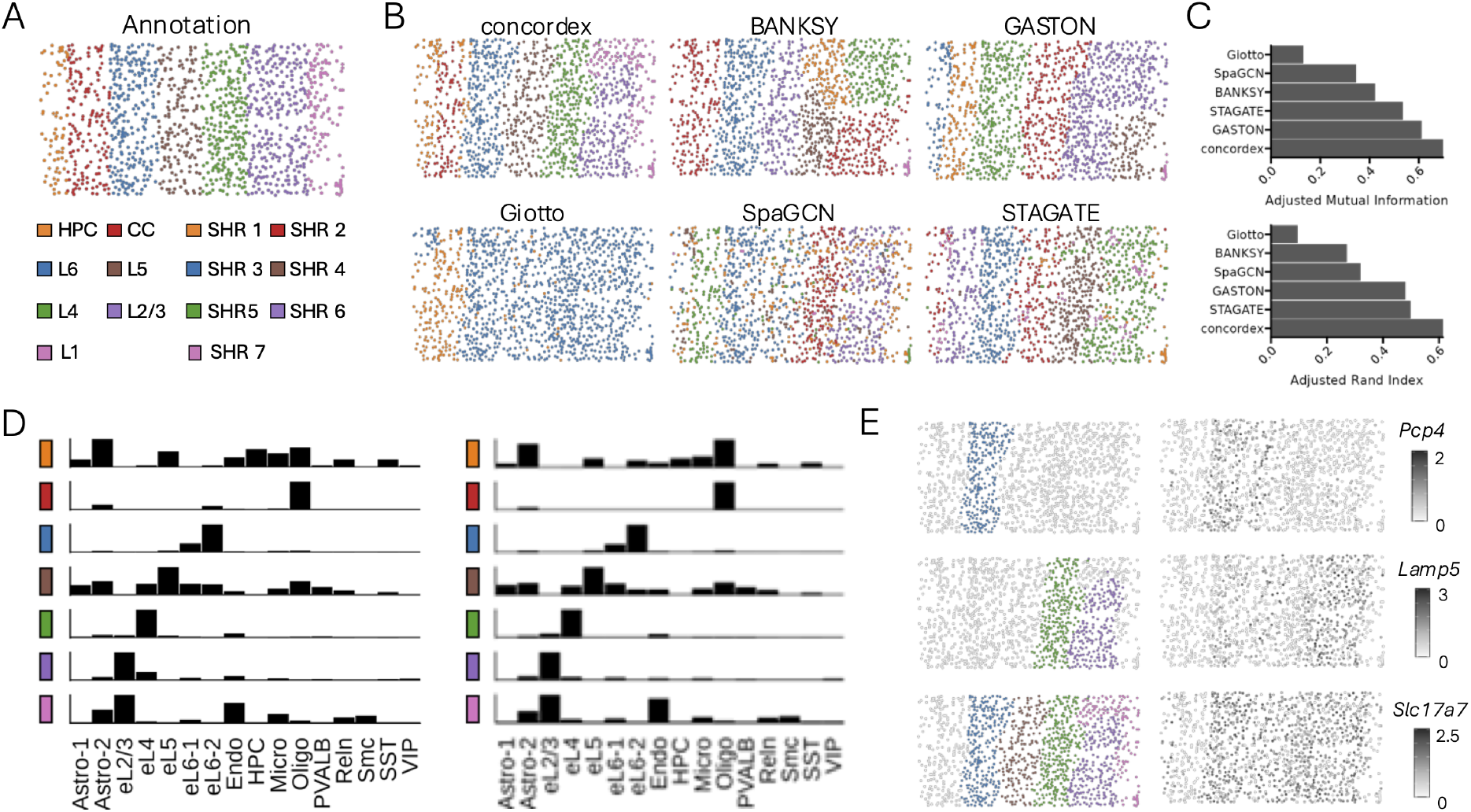
Evaluation of concordex on STARmap dataset. **A**. Manual annotations of the regions. **B**. SHR predictions from various methods. For concordex, the first 50 PCs were used to generate the neighborhood consolidation matrix. **C**. Performance of all methods using AMI (top) and ARI (bottom). **D**. Proportions of cell types in each SHR in the annotation (left) and concordex (right). **E**. Validation of predicted SHRs using normalized expression of known marker genes. *Abbreviations:* HPC - hippocampus, CC - corpus callosum

The combination of cellular and region annotations allows for a thorough evaluation of how well the regions predicted by concordex capture the expected cell type distribution in a well-studied system. The predicted SHRs agreed well with the annotations when run with default parameters (Figure 3B) and improved the predictions generated by other methods (Figure 3C). In contrast to Visium, the distance between adjacent spatial locations in the STARmap data varies considerably. The performance of concordex in this dataset also demonstrates that this approach is robust to differences in spatial scale that exist across SRT technologies. Additionally, the cell types present in each predicted SHR align with the expected cell type distributions in each layer even though specific cell types are present in multiple layers in similar proportions, for example, eL2/3 cells in L1 and L2/3 (Figure 3D). This is expected given that concordex explicitly clusters cells into regions on the basis of local neighborhood composition. Other methods fail to resolve this difference and combine the layers or produce arbitrary divisions (Figure 3B).

We further validated the predicted SHRs based on known marker genes for the upper and lower layers of the mouse neocortex (Figure 3E). The expression of *Slc17a7* is broadly expressed in all of the layers and excludes the white matter and hippocampus. To delineate the upper layers, we focused on *Lamp5* for layers 2/3 and 4. The expression of *Pcp4* has been shown to localize to the upper layers, L5 and L6. Importantly, the expression of these genes maps onto several SHRs predicted by concordex. This demonstrates the value of defining regions using rather than the spatial expression of individual genes.

### Identification of spatially variable genes in the mouse cerebellum

We next used concordex to identify SHRs in a Slide-Seq V2 mouse cerebellum dataset (22). Typical downstream differential expression (DE) analysis to identify spatially variable genes would examines differences between SHRs or rely on global metrics like Moran’s I or its local equivalent (23). However, we maintain that these approaches preclude exploration of the interaction between cell type and SHR, and cannot distinguish patterns that are driven by distinct spatial and cell type effects.

The SHRs identified by concordex show qualitative agreement with the annotations from the Allen Brain Reference Atlas (Figure 4A-B). To identify spatial DE genes, we used a Negative Binomial generalized linear model (NB-GLM) that includes an offset term for sequencing depth and an interaction term between cell type and SHR. In total, 1,246 genes could be explained by cell type, SHR, or an interaction (Figure 4C). We classified genes as being cell type- or spatial-dominant based on the model term with largest coefficient. For genes with evidence of an interaction, we further identified whether this effect could be explained more by cell type or SHR (Methods).

**Fig. 4.**
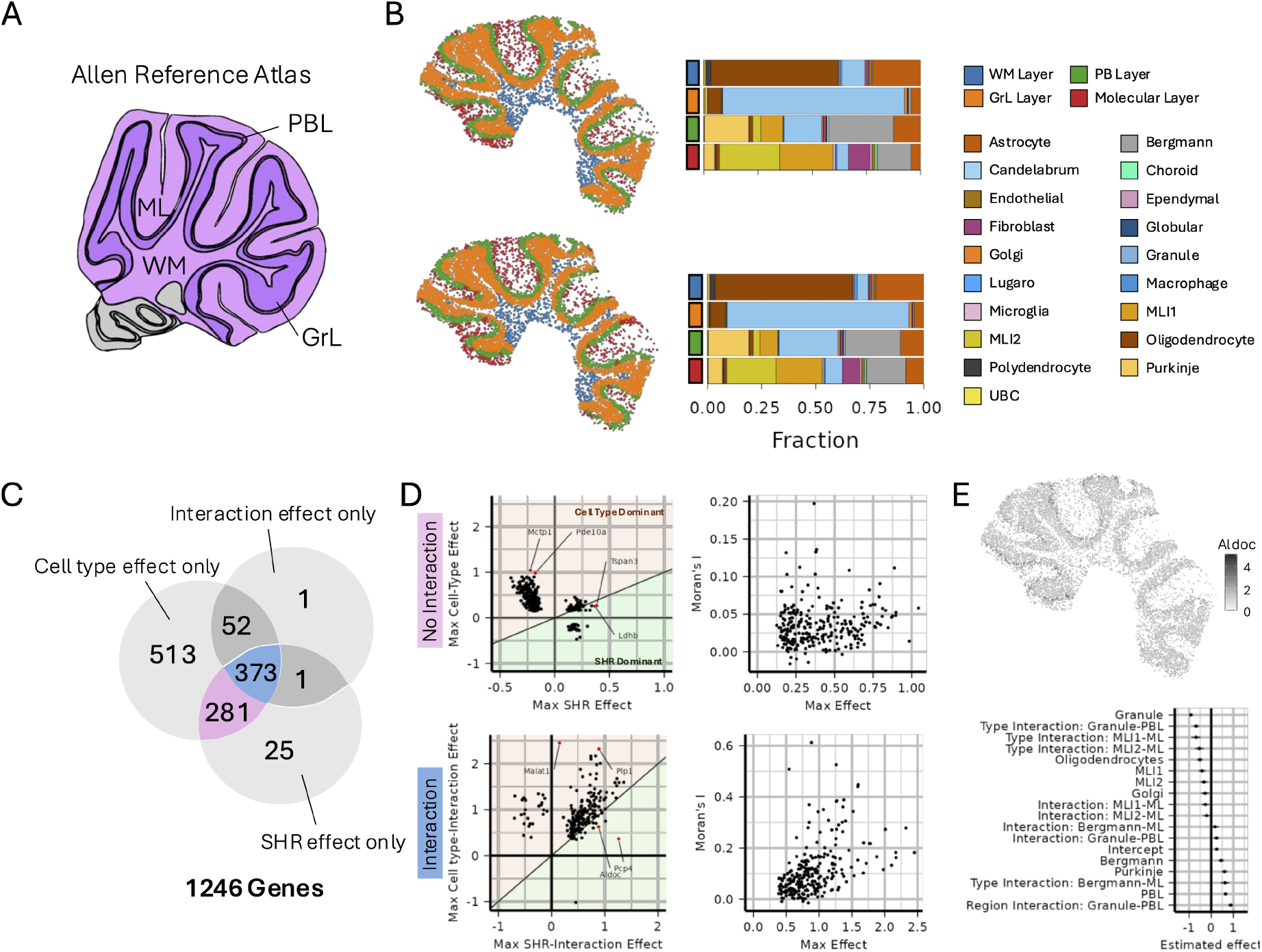
Identification of spatially variable genes using generalized linear models. **A**. Schematic of the mouse cerebellum from Allen Brain Atlas. **B**. Prediction of SHRs in the cerebellar cortex using cell types (top) or the top 50 PCs (bottom) to compute the neighborhood consolidation matrix. The proportion of cell types in each SHR is similar in either case.**C**. Different combinations of effects were detected in each gene. **D**. For genes without an interaction effect (top) cell type effects tend to be most prominent and correlate poorly with Moran’s I. In contrast, genes with an interaction effect (bottom) correlate more strongly with Moran’s I and are generally larger in magnitude. **E**. The gene *Aldoc* displays cell type and SHR-specific expression, the spatial effects dominate.

When we looked at genes that had a combination of SHR and cell type effects without an interaction, we found that cell type effects dominate for most genes, and in general, SHR effects were smaller in magnitude compared to cell type effects (Figure 4D; upper panel). Interestingly, genes without an interaction did not exhibit spatial autocorrelation measured by Moran’s I (Figure 4D; upper panel), which suggests that cell type and spatial interaction is required for autocorrelation. In contrast, genes with an interaction display strong spatial autocorrelation and have higher effects overall(Figure 4D; lower panel). The dominant effect for most genes in this class is the combined cell type and interaction effect. This suggests that the gene is more aptly described as a spatial-cell type marker gene. We observed that spatial-interaction effects are in general smaller than cell type-interaction effects (Figure 4D; lower panel) and in most cases, the spatial interaction effect is only slightly larger than the cell type interaction effect. One clear outlier to this pattern is *Aldoc* which has dominant spatial-interaction effects (Figure 4E). In the cerebellum, this gene is predominantly expressed by Purkinje cells, but it also exhibits strong spatial patterning, with expression localized to alternating parasagittal bands (24). The spatial banding pattern in Purkinje cells is apparent in the Slide-Seq data (Figure 4E), but the expression of this gene by other cells in this layer contribute to the strong spatial-interaction effect relative to the Purkinje-interaction effect (Figure 4E). Overall, these results show that linear models paired with accurate region identification by concordex can identify biologically and spatially relevant genes.

### Fast identification of SHRs in the mouse small intestine

Finally, we applied concordex to a VisiumHD dataset of a mouse small intestine (25). isiumHD greatly enhances the resolution of the Visium platform with the capture area 2*µ*m bins arranged in a regular 3250 × 3250 grid to achieve cellular scale. Nearly 400,000 8*µ*m bins overlap with the tissue sample in the mouse intestine dataset. Since this data is very sparse, we used the dataset that aggregated the 2*µ*m bins into the larger 16*µ*m bin size in order to visualize SHRs (Supplemental Figure S7). Using putative cell type annotations as input, concordex readily identified villus, crypt, and muscular structures that are consistent with histology (Supplemental Figure S7). These results demonstrate that concordex can accurately reconstruct spatial structures across diverse tissue types, even when working with large datasets, which are expected to become increasingly common in the future.

## Discussion

Efforts to characterize the expression and functional similarities of cells in their tissue context rely on accurate methods to identify regions with compositional similarity. We developed concordex to explicitly aggregate cells into regions based on the compositional similarity of their local neighborhoods. This approach enables long-range identification of regions and broad characterization of tissues. Our method is fast and flexible, leveraging research that has resulted in optimized algorithms for computing the kNN graph. On simulated data, concordex readily identifies global organization, even when the same cell types are represented throughout the spatial field. Using concordex, we were able to identify the well-described laminar structure of the mouse cerebellum and regions of functional importance in the mouse liver.

Importantly, we have demonstrated the utility of using local neighborhood compositional similarity as a marker of SHRs in spatially resolved transcriptomics data. Approaches that rely on k-NN graphs and discrete labels have been developed for multiplexed imaging analysis (14–16) and SRT (26). concordex extends these workflows both as a tool for exploratory analysis of non-spatial transcriptomics data and by allowing continuous attributes for neighborhood consolidation. We demonstrate that using continuous attributes to build SHRs performs similar to the discrete case. This addresses a significant limitation of other methods that to our knowledge, require cell type annotation as a prerequisite. Other methods aim to detect regions within a tissue where gene expression is consistent. The assumption is that the organization of tissues is related to the spatial dependence of gene expression. However, this approach for region identification often overlooks the cell type heterogeneity within a region and confounds the biological interpretation of spatial domains with the procedure used to generate them. For example, the notion of a ‘tissue domain’ in the BANKSY paper (8), is defined as the result obtained when ‘building aggregates with neighborhood kernel[s] and spatial yardstick[s]’. Similarly, in the GASTON paper (12), ‘spatial domains’ are described in terms of topographic maps, that result from isodepth which the GASTON method infers. Again, the notion of a ‘spatial’ or ‘tissue’ domain is tautological with the algorithm used to produce it. In the concordex framework, we prioritize the biological definition of spatial homogeneous regions, and our approach to identify SHRs follows from the definition, not the other way around. Thus, while other methods can, at times, produce similar results to concordex, concordex reliably distinguishes between cell type and region assignment and is particularly adept at identifying SHRs that recur in spatially distant parts of the tissue.

Many SRT studies aim to identify the relative position of cell types in space. Implicit in these analyses is that cell types are organized into SHRs, and efforts to identify region-specific variation largely rely on alignment to previously characterized anatomical structures (27). On the other hand, computational approaches for identifying SHRs vary in their scalability and interpretability. Spatial smoothing approaches often increase the dimension of SRT data, usually by concatenating information from spatial neighbors into a single matrix as input to dimension reduction and clustering algorithms (8, 28). These approaches are computationally burdensome as *k* (the number of neighbors) and *n* (the number of observations) becomes large, and can be intractable even for current datasets. As spatial transcriptomics technologies continue to improve, not only in terms of resolution, but also throughput, computational efficiency will become increasingly important.

Aggregation of cells and spots into neighborhoods based on the compositional similarity of their local neighborhoods enables long-range SHR identification of regions and broad characterization of the tissue. As we showed in simulation, existing methods that are based on gene expression or attributes derived from gene expression can fail to differentiate cell type variation from regional variation. By using information about cell neighborhoods, concordex naturally allows for cells of the same type to be assigned to different regions. When cell type labels are used with concordex, one possible limitation is that densely populated cells with identical neighborhoods may be identified as a single SHR, which may not correspond to a histological feature. We believe that this result is important for what it reveals about the tissue organization, such as varying cell density and type homogeneity.

We also demonstrated that concordex has non-spatial applications when the neighborhood consolidation matrix is constructed from principal components or expression vectors. A typical use case of concordex in this context includes assessing the existence of and relationships between predefined groups or clusters. In contrast to UMAP, the similarity matrix can be used to visualize distinct clusters without distorting the global relationships between them. The neighborhood consolidation matrix is especially useful for estimating the proximity of clusters and presents a more natural interpretation of the biological relationship between them. Given that UMAP plots are also used to visualize gene expression data within a cluster, we note that gene expression can be readily plotted as a heatmap grouped by pre-defined clusters without loss of information present in the UMAP visualization.

In summary, concordex provides an accurate and efficient framework for identifying SHRs across a variety of spatial scales and technologies, furthering the understanding of complex spatial regionalization patterns. Future work should focus on identifying genes with regionally restricted expression and distinguishing this pattern from cell type localization. We believe that the SHRs identified by concordex can offer substantial insight in future analyses and will facilitate further efforts to characterize complex tissues.

## Methods

### Overview of concordex

#### Construction of the kNN graph

The k-nearest neighbor (kNN) graph *G* = (*V, E*) can be generated from a scRNA-seq or SRT dataset where *V*, the set of cells or spots, and *E*, the set of edges, are determined according to some metric on *V*. The number of cells in the dataset is denoted |*V* | = *n*.

For SRT data, we use the Euclidean distance between the spatial locations of cell (or spot) *i* and cell *j* to determine the number of neighbors represented in the *G*, and disallow self-neighbors. By default, 30 nearest-neighbors are retained. For Visium data, we compute the 12 nearest neighbors.

#### Computation of the neighborhood consolidation matrix

The concordex workflow requires labeling the nodes of *G* from a finite set *C* with |*V* | = *m* distinct labels or with continuous vectors to create the *n* × *m* neighborhood consolidation matrix, *K*.

We conceptualize the case for continuous labels by considering the *k* nearest-neighbors of a node as a (*k −* 1)-simplex whose vertices can be described in ℝ^*m*^ for *m ≥ k*. When continuous vectors are used to color the nodes in *G*, this amounts to assigning those vectors to the vertices of the neighborhood simplex. This approach is easily amenable to discrete labels, where each discrete label is assigned to a unique element of a vector in ℝ^*m*^. The vertices can then be mapped to the standard vectors in ℝ^*m*^. In either the discrete or continuous case, the *n* rows of *K* can be interpreted as the center of mass of each simplex. We use the principal components to compute concordex in the DLPFC and STARmap datasets. However, alternative approaches such as Non-negative Matrix Factorization (NMF), topic modeling, or methods that identify gene programs in cells would also serve as equivalent options for this application. These methods generate latent representations that capture underlying patterns in the data that can be interpreted in a similar manner to PCA components. In the absence of explicit cell type information, these methods offer a way to abstract cell type into meaningful representations, effectively capturing its influence without requiring direct categorization. The columns of *K* either represent each of discrete labels or the non-zero elements of the vectors in ℝ^*m*^. For discrete labels, the entries *K*_*ij*_ can be interpreted further and represent the fraction of neighbors of cell *i* that are assigned label *j*.

#### Identification of spatial homogeneous regions

To assemble cells or spots into spatial homogeneous regions (SHRs), we use the matrix *K* and the k-means clustering algorithm when there is prior information for the expected number of domains. However, when the number of SHRs is not known, unsupervised clustering algorithms such as Leiden (17) can be used to generate assignments.

### Differential Expression Analysis

To assess whether spatial gene expression effects were a result of cell type effects, global spatial effects, or an interaction between the two, we used a Negative Binomial generalized linear model (NB-GLM) where the parameters were cell type, SHR, an interaction term, and an offset for library size. The link function for the mean can be written as

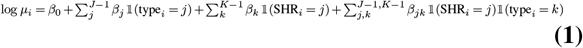

In the equation above, 𝟙_(*·*) is the indicator function and the *β* coefficients are estimated from the model. The linear models were fitted and analyzed using the INLA R package (29).

To classify genes by the pattern of cell type and spatial effects, we first excluded any terms with a confidence interval including zero. After this exclusion, terms without evidence of an interaction were called “cell type dominant” if a cell type term had the greatest effect and were called “spatial dominant” if a spatial term had the greatest effect. For genes with non-zero interaction terms, we first rearranged equation 1 to group like spatial or like cell type terms. In the former case, we computed total effect of SHR *k* interacting with cell type *j* as *β*_*k*_ + *β*_*jk*_. Similarly, we computed the total effect of cell type *j* interacting with SHR *k* as + *β*_*jk*_. We refer to these effects as spatial-interaction and cell type-interaction effects, respectively. Similar to the non-interaction case, genes with a greater spatial-interaction term were called “spatial-interaction dominant”, otherwise, we labeled the gene “cell type-interaction” dominant. In cases where the fixed cell type or SHR effect was greater than the interaction effects, we labeled the gene “cell type dominant” or “spatial dominant”. Genes with only negative terms were not classified.

### Datasets

We applied concordex to samples from several SRT technologies including Visium, VisiumHD, STARmap, and Slide-Seq V2. More specifically, we used the human DLPFC Visium dataset from (19). Each of the 12 samples in this dataset contained approximately 3460 to 4789 spots and profiled more than 12,000 genes. Manual annotations of the cortical layers were provided with the data and served as ground truth in our analysis. We downloaded the mouse intestine VisiumHD dataset from the 10x genomics website (25). The data were provided at the 2*µ*m, 8*µ*m, and 16*µ*m resolution along with putative cell type assignments for the 8*µ*m and 16*µ*m resolution data. The STARmap data (21) contained 1020 genes measured in 1207 cells, and the Slide-Seq V2 (22) data contained 10,121 genes and 9985 cells. The details for the simulated dataset are provided in the Supplementary Information. Briefly, we used the splatter simulation software (30) to generate a count matrix containing 14,400 cells and 10,000 genes and arranged these cells in either a checkerboard or stripe pattern in different proportions according to the simulated cell type.

## Data pre-processing

In all datasets that rely on a patterned capture grid, we first removed all capture locations that were outside of the main tissue area. If raw data were available, genes expressed in fewer than 50 cells/spots were removed. We used the analytic Pearson residuals to normalize the data, compute principal components, and identify the top 3,000 highly variable genes. Otherwise, we used the processed data provided by the authors.

## Ethics approval and consent to participate

Not applicable

## Consent for publication

Not applicable

## Data availability

The data used in this manuscript are available from their original authors.

The Visium DLPFC dataset and annotations (19) are available for download using the SpatialLIBD Bio-conductor package here https://doi.org/doi:10.18129/B9.bioc.spatialLIBD. The STARMAP dataset from (21) is available on Figshare with the identifier https://doi.org/10.6084/m9.figshare. 22565209.v1. We obtained the Mouse Cerebellum Slide-Seq V2 dataset from (22) and downloaded it from the Broad Institute Single Cell Portal https://singlecell.broadinstitute.org/single_cell/study/SCP948. We downloaded the mouse intestine VisiumHD dataset from the 10x genomics website (25)

## Code availability

concordex is available as a Bioconductor package at https://www.bioconductor.org/packages/release/bioc/html/concordexR.html. The source code for the Bioconductor implementation is available at https://github.com/pachterlab/concordexR, and the Python version is available at https://github.com/pachterlab/concordex.

Code to reproduce the figures in this manuscript are available at the following Github repository: https://github.com/pachterlab/JBMMCKP_2023/

## Author contributions

The concordex project was led by KJ who conceptualized the spatial concordex method, obtained the results and performed the analyses in the paper, implemented the method in R, and drafted the manuscript. ASB and AGM implemented an initial version of concordex in Python, and contributed to the development of the non-spatial concordex method and applications. LM and TC helped in developing the concordex non-spatial method. AK improved the concordex Python implementation under the supervision of KJ. OHM contributed to the DE analysis. KJ and LP edited the manuscript with assistance from the other authors.

## Acknowledgments

The authors thank Michal Polonsky and other participants in the Caltech CI2 initiative for helpful discussions on challenges in interpreting spatial transcriptomics data.

