## Supplemental Information for "Characterization of spatial homogeneous regions in tissues with concordex"

### Supplementary Information Characterization of spatial homogeneous regions in tissues with concordex

#### Supplementary Note 1: Additional Methods

**Availability of analyzed datasets.** Here, we provide additional information for each of the analyzed datasets:

**Simulated datasets** For the checkerboard simulation in Figure **??** we created a square lattice with 120 rows and columns. Each individual checker box had a dimension of 15 rows and columns so that the grid resembled a traditional checkerboard. Each square in the lattice represents a 'cell' for a total of 14,400 cells.

We used the splatter simulation software (1) to generate a count matrix containing 14,400 cells and 10,000 genes. We updated the 'group.prob' parameter to generate two cell types with distinct gene expression profiles. All other parameters were kept at their default settings. A total of [number] cells were labeled 'Type 1' and the remaining cells were labeled 'Type 2'. In the white regions of the checkerboard, we assigned 80% of the squares 'Type 1' and in the black regions, 20% of the squares were assigned 'Type 1'.

For the results in Figure S6, we generated a laminar pattern in the grid by grouping squares every 30 columns to create 4 layers. In the layered configuration, each stripe had approximately 25%, 40%, 60%, and 75% of the locations within the group assigned to 'Type 1', respectively.

**Slide-Seq V2** We downloaded the processed count matrix, spatial locations, and cell-type labels for Replica 1 from reference (2). The dataset contains counts for 11,626 spatial locations and 23,096 genes. The pixels were filtered to include only spots with a minimum of 100 UMIs detected and those that were not labeled 'reject'. This left a total of 9,985 pixels available for analysis. A detailed description of the preprocessing steps is available (2).

**STARmap** We obtained a copy of the STARmap dataset from Figshare at (3). The dataset included counts for 1020 genes measured in 1207 cells.

**Visium** We downloaded the count matrices, spatial coordinates, H&E images, and spot annotations from the original authors (4). Each of the 12 samples in this dataset contained 3460 to 4789 spots that overlapped with the tissue slice and profiled more than 12,000 genes.

**VisiumHD** The Visium HD Mouse Small Intestine (FFPE) dataset can be downloaded from the 10x Genomics data download portal (5). We included all bins that overlapped with the tissue slice in the analysis.

**10x Chromium mouse ex-utero embryo** The datasets used in this study were derived from the reference (6). A detailed description of the preprocessing steps are described elsewhere (7). Briefly, the authors provided two transformed count matrices. The 'Log-Normalized' dataset consisted of log-transformed counts, and the 'Stabilized-Scaled' dataset had been variance stabilized using Seurat and subsequently mean-centered and scaled. Both datasets were integrated using Seurat. The count matrices were mean-centered and scaled before principal component analysis (PCA). PCA analysis was performed using sklearn TruncatedSVD to 15 dimensions to agree with the analysis performed by the original authors. We used the findkNN function from the BiocNeighbors package in R to find the exact 30 nearest neighbors for each cell in the dataset.

The UMAP algorithm was applied to the 15-dimensional PCA embeddings with default settings except where noted. The subsequent visualizations were colored using labels provided by the authors.

**Benchmarking concordex with other methods.** In the section below, we describe how we ran each of the competing methods highlighted in the main text:

**BANKSY** We followed the examples in the package documentation and used the author's guidance to set parameters. Specifically, we used 'lambda=0.8' for spatial domain detection in all comparisons and only used the top 3,000 HVGs in the workflow.

**BayesSpace** BayesSpace provides methods for clustering SRT data obtained on the Visium, VisiumHD, or Spatial Transcriptomics platforms. We applied BayesSpaces to the DLPFC dataset following the example here: `http://www.ezstatconsulting.com/BayesSpace/index.html`. Except for the number of clusters, all other parameters to the 'spatialCluster()' function were assigned to the default.

**GASTON** We ran GASTON according to the tutorials available on the software Github (`https://github.com/raphael-group/GASTON`). Running GASTON requires training a neural network with different initializations and choosing the model with the lowest loss. We used 15 initializations for each of the datasets. We used the network architecture suggested by the authors in the mouse cerebellum tutorial. When available, we provided the number of desired spatial regions and set all other parameters to their defaults.

**Giotto** To run Giotto, we followed the 'Spatial Domain Analysis using HMRF' vignette found here: `https://drieslab.github.io/Giotto_website/articles/hmrf.html`. First, to identify spatially variable genes, we ran the 'binSpect()' function with default parameters and selected the top 500 genes for input to the 'doHMRF()' function.

**STAGATE** We followed the author's guidance to run STAGATE on various processed datasets (`https://stagate.readthedocs.io`). As noted in the manuscript, we used the analytic pearson residuals to normalize the datasets in this study rather than depth normalization. Downstream parameters were set according to the author vignettes, if available, and left at the defaults otherwise.

**SpaGCN** For all analyses, we followed the step-by-step tutorial on the Github (`ttps://github.com/jianhuupenn/SpaGCN`). We set the hyperparameter 'p=0.5' for all datasets and all other parameters were left to their defaults. As with other methods, we used the analytic Pearson residuals to normalize the datasets in this study instead of depth normalization.

#### Supplementary Note 2: Additional Applications

**Using concordex to Evaluate Cell type Clustering and Integration Effects.** We evaluated the ability of concordex to assess the fidelity of cell type clustering and dataset integration on data generated from in-utero and ex-utero mouse embryos at the E10.5 developmental stage (6). We evaluated the data in the 15-dimensional PCA space and 2-dimensional UMAP embedding by computing the neighborhood consolidation and similarity matrices using cell type or growth condition as the label.

Using cell type labels, we found that the concordex similarity matrix can readily visualize distinct clusters in both the PCA and UMAP embeddings. In the log-normalized PCA data, concordex suggests an relationship between some cell-type clusters (Supplementary Figure 3A). For example, the neighborhoods of cells in cluster 0 are composed of cells from clusters 5, 7, 10, and 13. We observed that simply changing the random state improves the qualitative appearance of the cluster separation in the UMAP embedding, reflecting the arbitrary nature of otherwise equivalent embeddings. While tuning the hyperparameters can alter how well the UMAP embedding preserves the structure of the PCA space, concordex directly visualizes the cluster relationships in the PCA-embedded data and reduces the need to embed the data in fewer dimensions.

Consistent with (7), we observed that the UMAP embedding can be a poor representation of the kNN graph in general. For example, inspection of the clusters in the scaled-stabilized data UMAP embedding indicates a close relationship between several clusters even though this structure is not apparent in the kNN graph generated from the PCA-embedded data (Supplementary Figure 3B). The neighborhood consolidation matrix allows for the assessment of the degree to which clusters overlap; e.g., it can show which cluster has the greatest number of cells where more than half of their neighbors have a different label. We found that the number of cells with mixed neighborhoods is often greater in the UMAP embedding (Supplementary Figure 4), showing that clusters are more mixed in the UMAP embedding than they are in the PCA space. These relationships are especially difficult to glean from the UMAP embedding alone, where overlapping clusters can appear indistinguishable, whereas concordex provides an exact assessment of the extent of mixing between clusters. Moreover, the concordex similarity matrix is sufficient to visualize the global structure of the kNN graph when combined with the concordex measures.

**Architecture of the mouse cerebellum.** We also applied the method to mouse cerebellum data that was generated using Slide-Seq V2 (8). Though this method has near cellular resolution ($10\mu$m), spatial capture spots can contain information from more than one cell. We therefore used the cell type labels that were published with the original manuscript and determined by the spot deconvolution method RCTD (2). This tissue is known to contain well-defined domains (2) namely the molecular, Purkinje-Bergmann, Granule, and white matter layers. Since these regions also have distinct cell type composition, we reasoned that concordex should be able to detect the boundaries between each layer.

Inspection of the neighborhood consolidation matrix highlights that SHRs do not need to be dominated by a single cell type (Figure S2A). Qualitative assessment of the predicted SHRs show agreement with the expected morphology of the tissue and contain cell types in proportions similar to canonical expectations (Figure S2B). Notably, the performance of concordex is comparable to more complex methods that rely on neural networks (Supplementary Figure S1). In each SHR, we compared the prediction to the RCTD weights of the most dominant cell types in the region (Figure S2D). All regions predicted by concordex correspond strongly to the cell type localization, even in areas of low cell density.

The SHRs predicted by concordex share the most agreement with the regions predicted by GASTON (Supplementary Figure S1). Visual inspection of the regions predicted by STAGATE and SpaGCN visually resemble the expected arrangement in the cerebellum, but inspection of the cell type proportions in each region reveals a failure to resolve the canonical layers (Supplementary Figure S1). In particular, STAGATE does not distinguish between the molecular and Purkinje layers. This is most evident in the bar graphs showing the cell type composition of each layer where there is little difference between the composition predicted Purkinje and molecular cell layers.

To demonstrate that concordex can produce meaningful results when discrete labels are not available, we used the top 50 PC loadings to label each spot. We averaged the loadings of the neighbors of each cell to create the neighborhood consolidation matrix and used this as input to clustering algorithms as above. Using this labelling approach, each row of the matrix is a spot, each column is a PC, and the entries are the average loadings of the PCs across neighbors. The resulting SHRs are qualitatively similar to the results using discrete labels (Figure S2C) and most spots receive the same assignment with either labeling scheme.

Since concordex should only aggregate spots into a SHR if they share similar neighborhood composition, we tested whether concordex could detect manipulations to the cell type identity of a subset of spots. We chose a subset of spots in the molecular layer and randomly reassigned their cellular identity by swapping cell labels (Supplementary Figure S2). This created an artificial, heterogeneous region that was distinct from the organized layers in the remaining tissue (Supplementary Figure S2). We observed that concordex was able to distinguish the heterogeneous region from the remaining molecular layer and the organization of the unaltered tissue remained largely unchanged. Notably, this alteration significantly altered the regions predicted by GASTON (Supplementary Figure S2), resulting in an organization that did not correspond to the known tissue structure. Altogether, these results reveal that concordex is sensitive to varying cell type composition and can reliably distinguish regional differences in a tissue.

### Supplementary Figures

A. Log Normalized, Integrated Counts

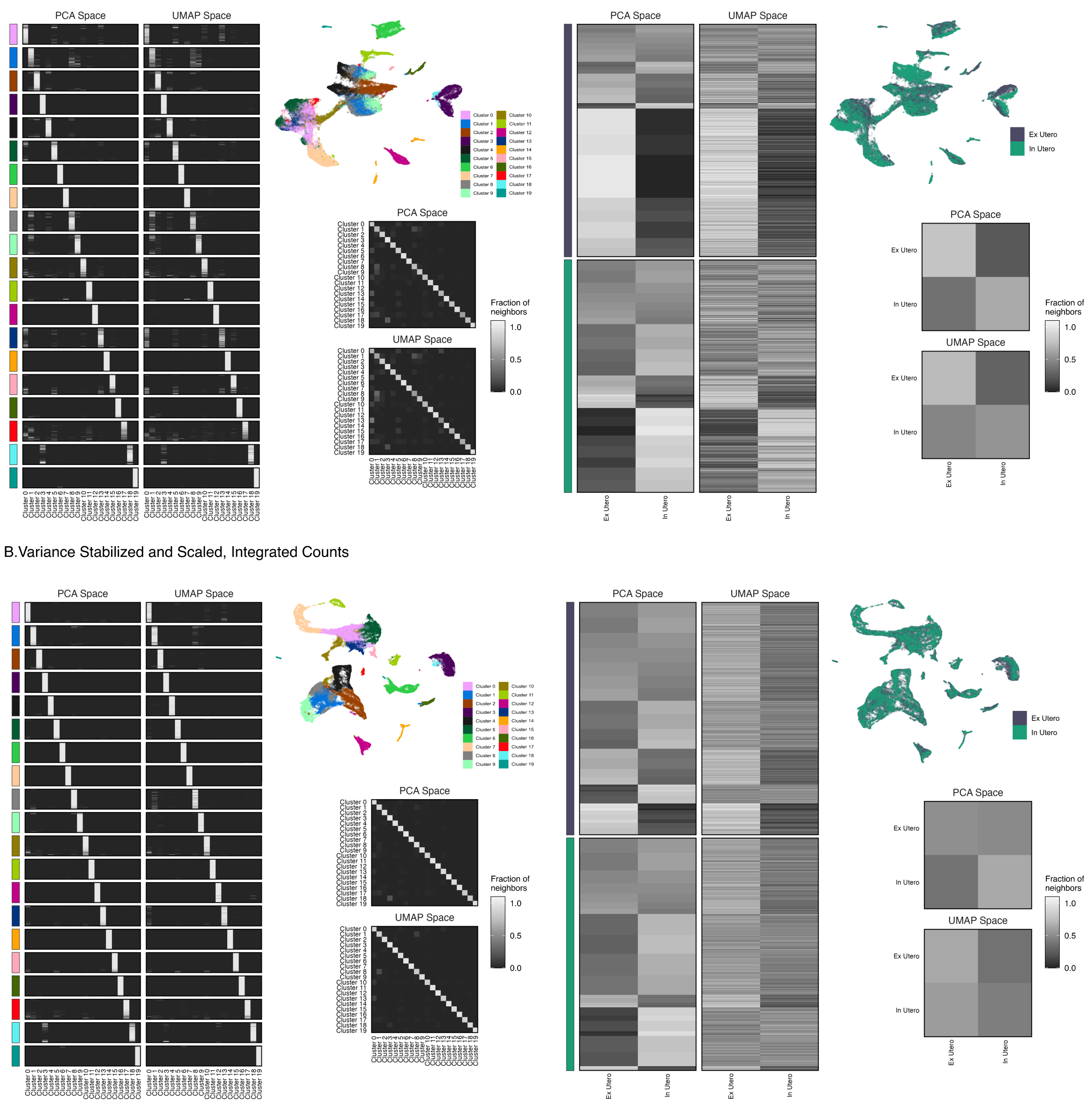


**Fig. S1.** Evaluation of concordex on mouse embryo generated from 10X Chromium V3. Left plots show the neighborhood consolidation matrices of the A) Log-normalized and integrated counts or B) Variance stabilized, scaled, and integrated counts with the UMAP embeddings and averaged similarity matrices next to it. In the neighborhood similarity matrix, each row is a cell and cells are grouped by their assigned cluster or batch. When the heatmaps are grouped by cluster assignment in the similarity matrices, the visualizations show that almost all clusters are well separated which contradicts visual inspection of the UMAP embedding. Similarly, visual inspection of the UMAP colored by growth condition suggests that the data are effectively integrated, however, the heatmaps show that most cells have neighborhoods that are exclusive of one growth condition.

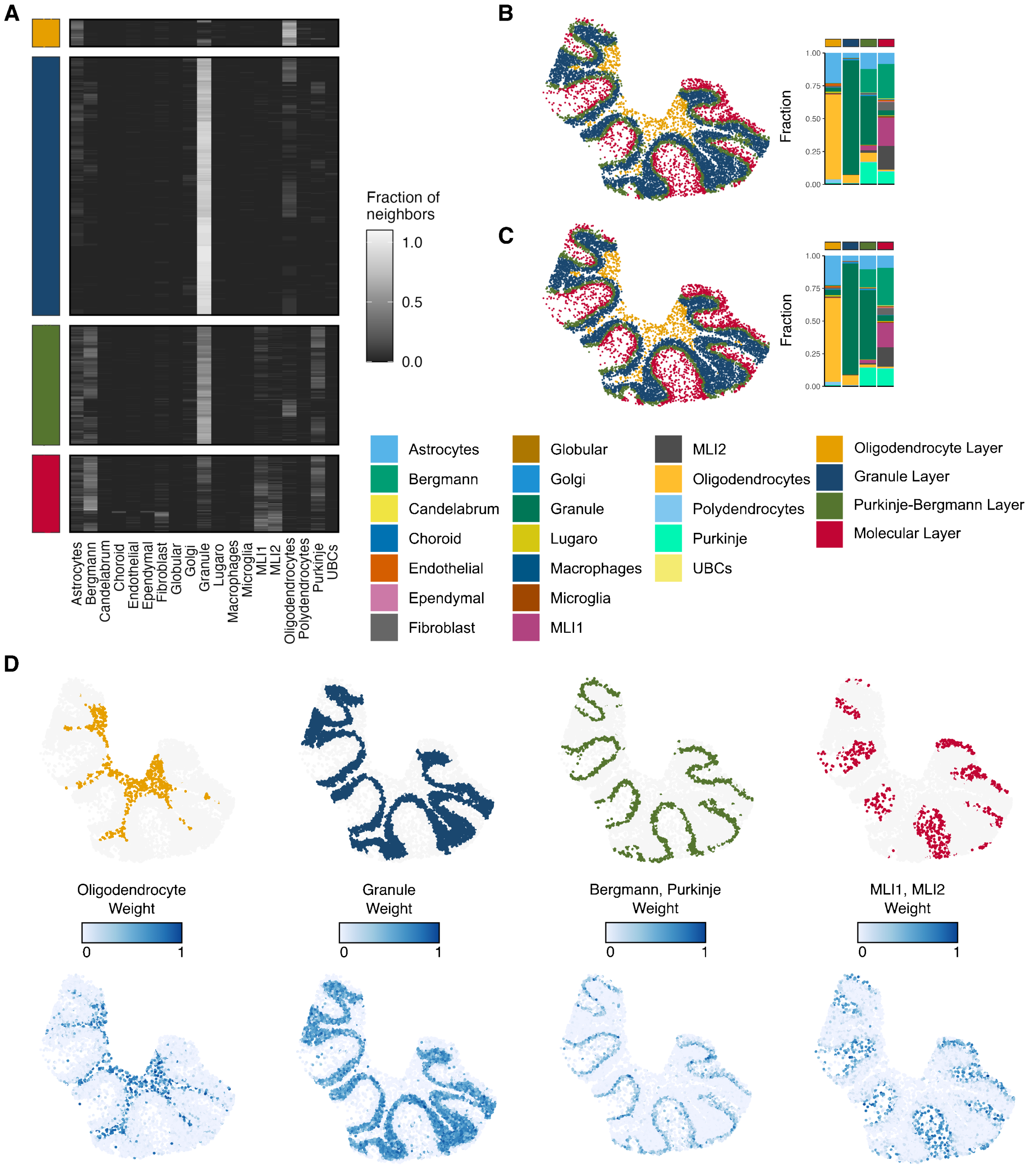


**Fig. S2.** Results of concordex on a slice of mouse cerebellum. A) A heatmap showing for each cell in each region, the fraction of neighbors of each cell types. B) Identification of spatial homogeneous regions with concordex based on cell type annotations. C) Identification of spatial homogeneous regions with concordex based on 50-dimensional PCA coordinates. D) Comparison of RCTD weights to distinct concordex spatial homogeneous regions.

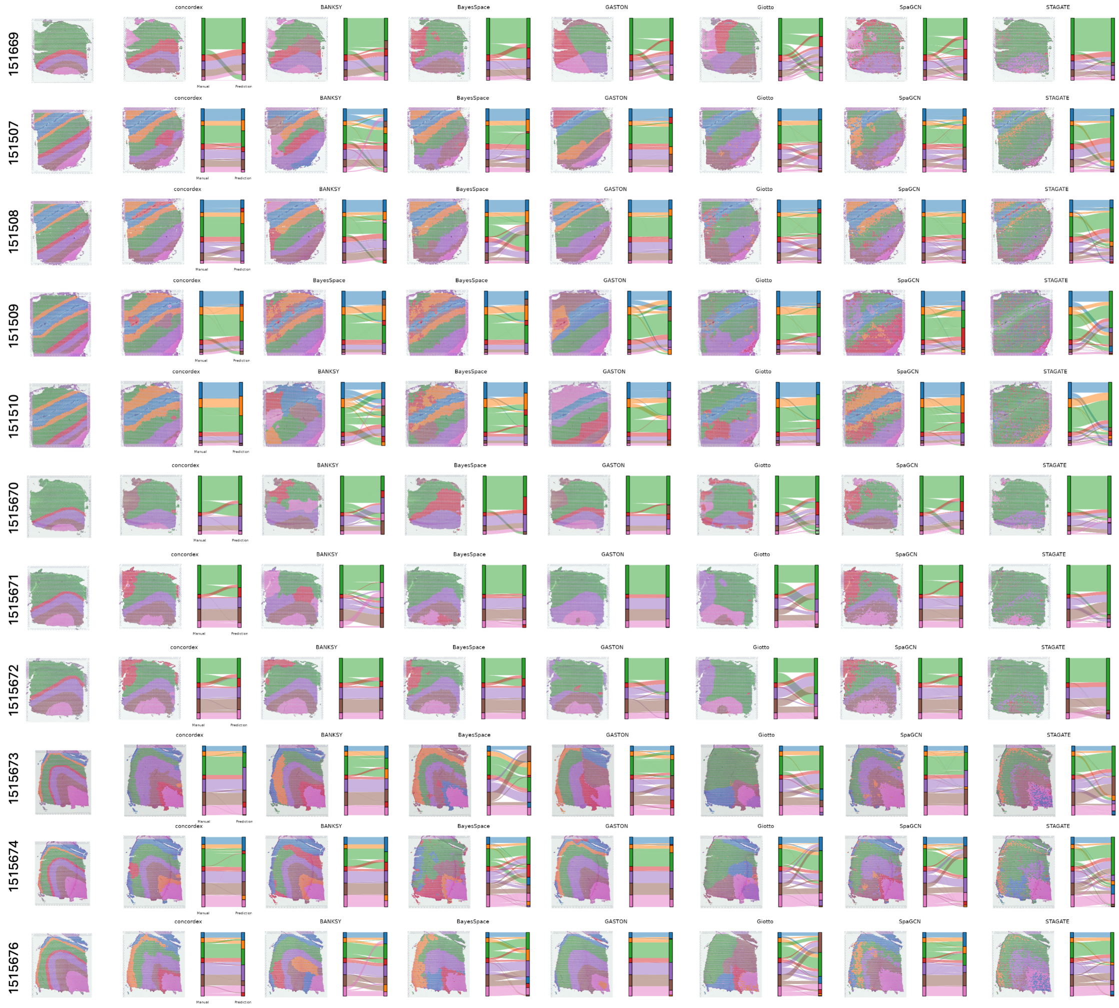


**Fig. S3.** Comparison of concordex to other methods on DLPFC visium dataset. For all samples, each panel shows manual annotations provided by the authors (left) and concordex results (right). The smaller insets are the results produced by BANKSY, BayesSpace, GASTON, Giotto, SpaGCN, and STAGATE.

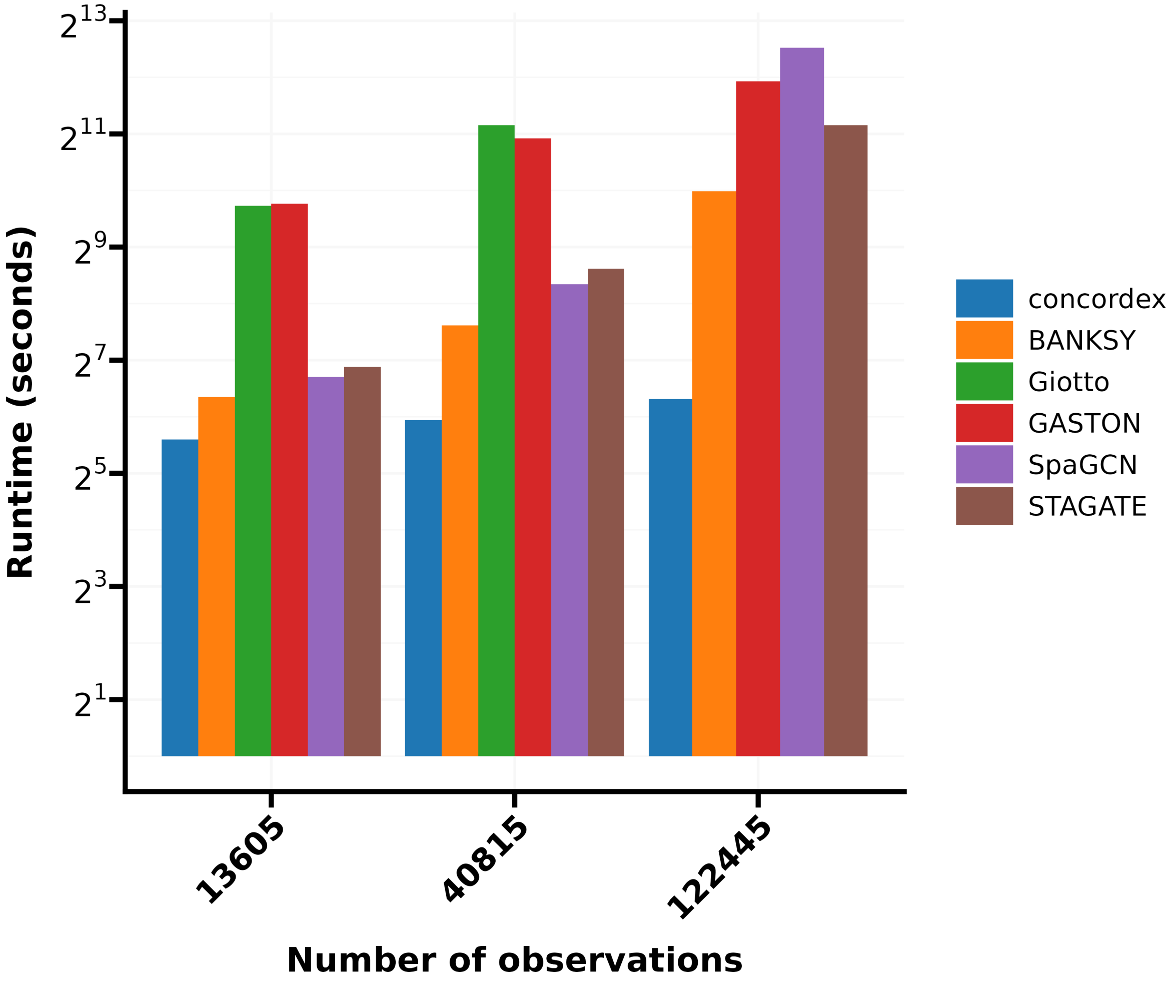


**Fig. S4.** Comparison of speed performance across all tested methods in a MERFISH dataset downsampled to the number of cells indicated on the x-axis.

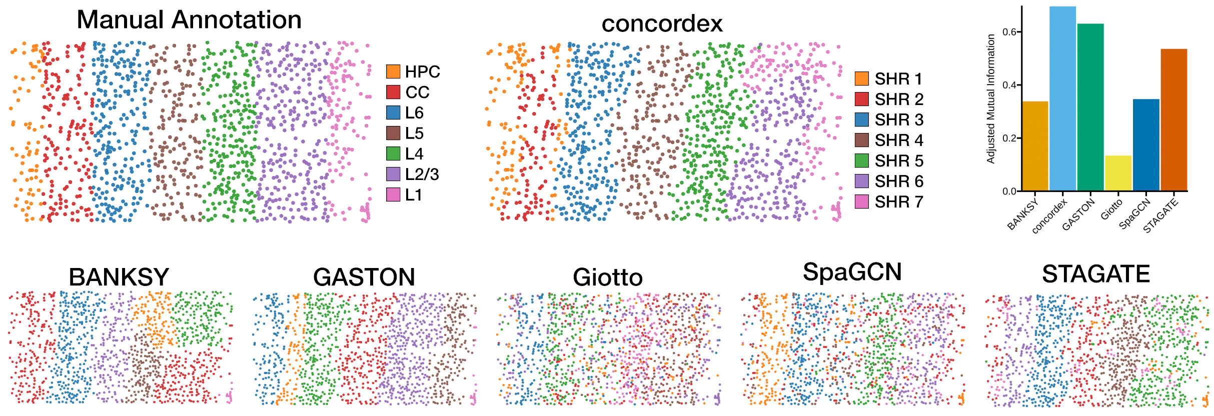


**Fig. S5.** Comparison of concordex to other methods on STARmap mouse V1 dataset. (A) Manual annotations of the regions identified in the mouse primary visiual cortex and (B) Results from concordex as in **??**. (C) Comparison of concordex to other methods via Adjusted mutual information. The results from other methods are shown in (D-H).

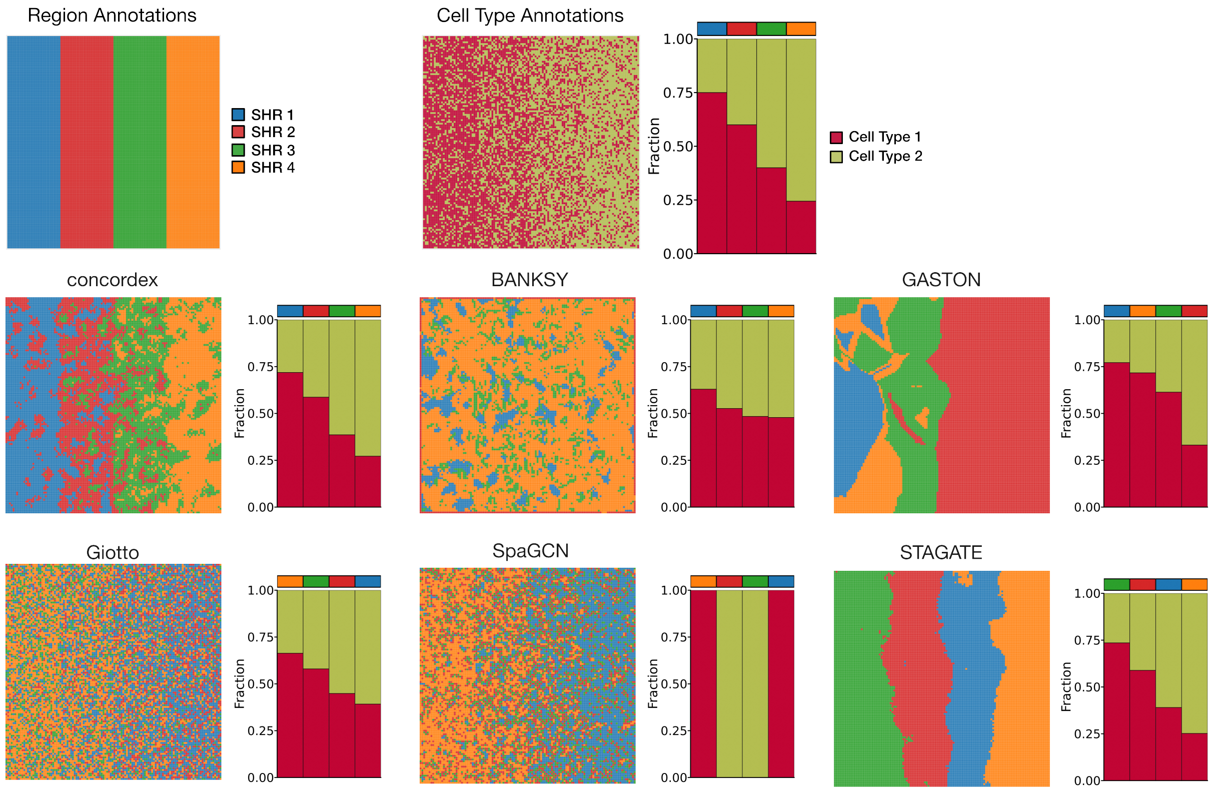


**Fig. S6.** Performance of concordex on synthetic data. (A-B) We designed a control experiment consisting of a gradient of regions, each with two cell types, and with increasing proportion of one cell type across the gradient. For concordex (C) we constructed the neighborhood consolidation matrix using the top 50 principal components. All other methods (D-H) were run as in ??

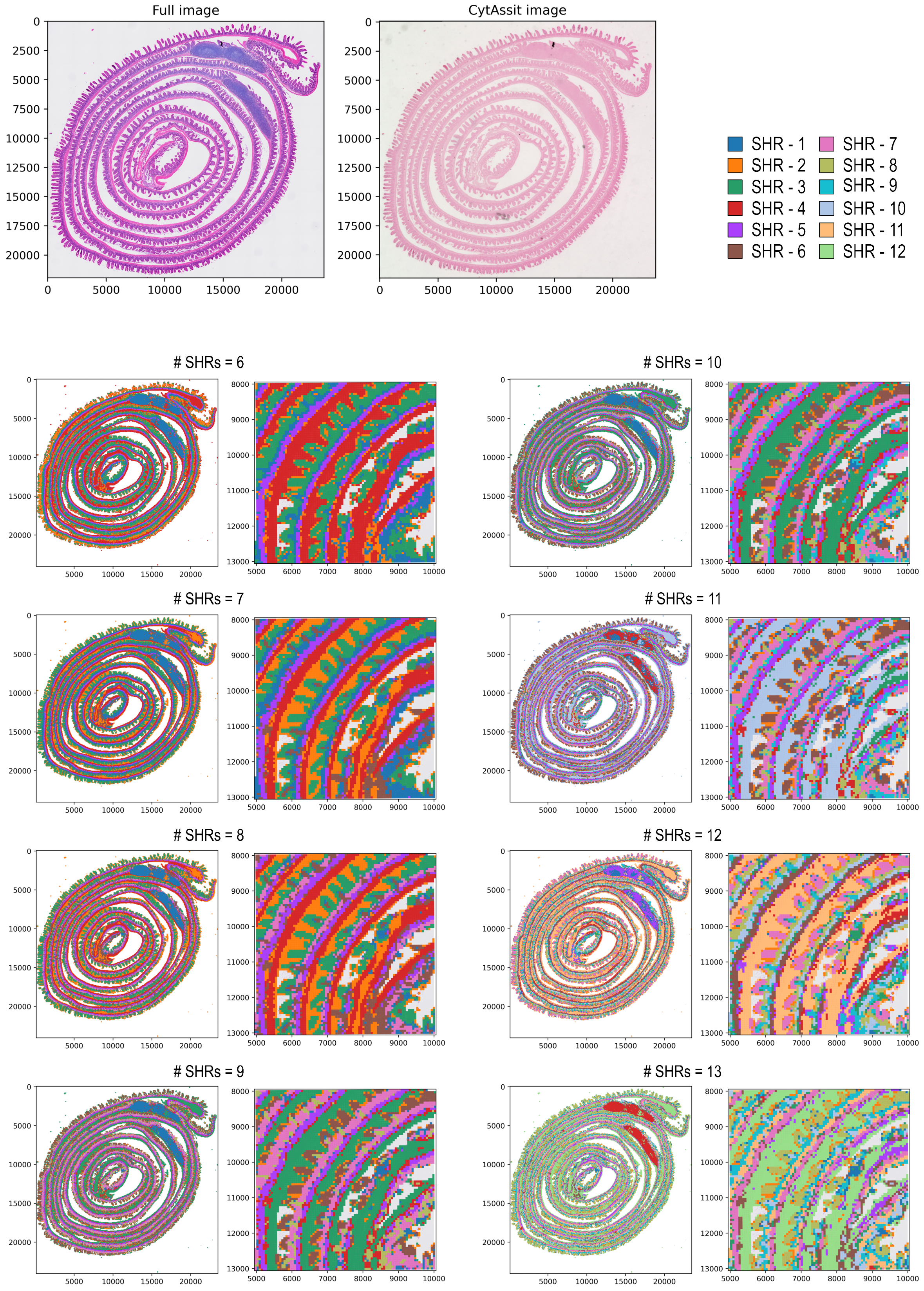


**Fig. S7.** Evaluation of concordex on Visium HD dataset. We applied cocnordex to the VisiumHD dataset of the mouse small intestine (A-B) and identified up to 13 SHRs (C-J) using the neighborhood consolidation matrix computed from cell type labels.
